# *Coxiella burnetii* Type 4B Secretion System-dependent manipulation of endolysosomal maturation is required for bacterial growth

**DOI:** 10.1101/645382

**Authors:** Dhritiman Samanta, Tatiana M. Clemente, Stacey D. Gilk

## Abstract

Upon host cell infection, the obligate intracellular bacterium *C. burnetii* resides and multiplies within the *<u>C</u>oxiella*–<u>C</u>ontaining <u>V</u>acuole (CCV). The nascent CCV progresses through the endosomal maturation pathway into a phagolysosome, acquiring lysosomal markers as well as acidic pH and active proteases and hydrolases. Approximately 24-48 hours post infection, heterotypic fusion between the CCV and host endosomes/lysosomes leads to CCV expansion and subsequent bacterial replication in the mature CCV. Initial CCV acidification is required to activate *C. burnetii* metabolism and the Type 4B Secretion System (T4BSS), which secretes effector proteins required for CCV maturation. However, we recently found that the mature CCV is less acidic (pH~5.2) than lysosomes (pH~4.8). Further, CCV acidification to pH~4.8 causes *C. burnetii* lysis, suggesting *C. burnetii* actively regulates CCV pH. Because heterotypic fusion with host endosomes/lysosomes may influence CCV pH, we investigated endosomal maturation in cells infected with wildtype (WT) or T4BSS mutant (Δ*dotA*) *C. burnetii*. We observed significantly fewer LAMP1-positive lysosomes, along with less acidic “mature” endosomes (pH~5.8), in WT-infected cells, compared to mock or Δ*dotA*-infected cells. Further, while endosomes progressively acidified from the periphery (pH~5.5) to the perinuclear area (pH~4.7) in both mock and Δ*dotA*-infected cells, endosomes did not acidify beyond pH~5.2 in WT-infected cells, indicating that the *C. burnetii* T4BSS inhibits endosomal maturation. Finally, increasing the number of acidic lysosomes by overexpressing the transcription factor EB inhibited *C. burnetii* growth, indicating lysosomes are detrimental to *C. burnetii*. Overall, our data suggest that *C. burnetii* regulates CCV pH, possibly by reducing the number of host lysosomes available for heterotypic fusion.

**Author summary:** The obligate intracellular bacterium *Coxiella burnetii* causes human Q fever, which manifests as a flu-like illness but can develop into a life-threatening and difficult to treat endocarditis. *C. burnetii,* in contrast to many other intracellular bacteria, thrives within a lysosome-like vacuole in host cells. However, we previously found that the *C. burnetii* vacuole is not as acidic as lysosomes and increased acidification kills the bacteria, suggesting that *C. burnetii* regulates the pH of its vacuole. Here, we discovered that *C. burnetii* blocks endosomal maturation and acidification during host cell infection, resulting in fewer lysosomes in the host cell. Moreover, increasing lysosomes in the host cells blocked *C. burnetii* growth. Together, our study suggests that *C. burnetii* regulates vacuole acidity and blocks endosomal acidification in order to produce a permissive intracellular niche.

## Introduction

*Coxiella burnetii* is a gram-negative obligate intracellular bacterium which causes human Q fever. Q fever manifests as a flu-like illness in acute disease and can develop into culture-negative endocarditis in chronic cases. The current treatment regimen for chronic *C. burnetii* infection requires a daily antibiotic combination therapy for at least 18 months [1], highlighting the need for more efficient therapeutics. Transmitted through aerosols, the bacteria are phagocytosed by alveolar macrophages and initially reside in a tight-fitting nascent phagosome that matures through the canonical host endocytic pathway. As early as 40 minutes post infection, Rab5 and Rab7, markers of early and late endosomes, are sequentially recruited to the *C. burnetii-* phagosome membrane [2]. Approximately 2-6 hours post infection, the phagosome fuses with lysosomes [3, 4], delivering lysosomal membrane proteins, including LAMP1 (Lysosome-associated membrane glycoprotein-1) and v-ATPase [5], and lysosomal enzymes such as acid phosphatases [4, 5] and cathepsin D [2, 5, 6] to the phagosome. Approximately 24-48 hours post infection, phagosome expansion, presumably through fusion with the endocytic pathway, gives rise to a large acidic phagolysosomal-like compartment termed the *Coxiella-*containing vacuole (CCV) [5].

*C. burnetii* effector proteins, which are secreted into the host cell cytoplasm through a Dot/Icm Type 4B Secretion System (T4BSS) [7, 8], manipulate host cell processes to support CCV expansion and bacterial growth [9–12]. Inhibiting *C. burnetii* protein synthesis by chloramphenicol treatment or inactivating the *C. burnetii* T4BSS results in smaller CCVs [9, 13], implicating T4BSS effector proteins in CCV expansion and subsequent bacterial growth. Interestingly, in the absence of *C. burnetii* protein synthesis the nascent phagosomes still acidified and acquired LAMP1, yet did not expand to become mature CCVs [13], suggesting that while early phagosome-lysosome fusion and acidification are not *C. burnetii-*dependent, *C. burnetii* regulate CCV expansion and maintenance. Early studies using fluorescein isothiocyanate (FITC) as a pH probe suggested that the CCV has an acidic pH similar to lysosomes (pH~4.5) [14, 15]. Further, acidic pH activates *C. burnetii* metabolism and T4BSS [16, 17]. Therefore, in contrast to many other intracellular bacteria which block phagosome-lysosome fusion, including *Legionella pneumophila, Mycobacterium tuberculosis, Anaplasma sp.,* and *Yersinia pestis* [18–22], *C. burnetii* survives in the phagolysosomal environment. We recently developed a ratiometric microscopy-based method to measure CCV pH using Oregon Green 488, a pH-sensitive fluorophore [23] and determined the CCV pH to be ~5.2 in both HeLa cells and cholesterol-free mouse embryonic fibroblasts (MEFs) [24]. In agreement with our results, a study with Chinese hamster ovary (CHO) cells measured pH of intact CCVs to be ~5.2 [25]. Moreover, we found that inducing CCV acidification to pH ~4.8 through cholesterol accumulation in the CCV membrane led to bacterial lysis [24]. This surprising finding suggests that *C. burnetii* is sensitive to the more acidic pH of lysosomes, and led us to hypothesize that, in contrast to previous results, *C. burnetii* does indeed regulate the pH of the intracellular niche.

The CCV is highly fusogenic and acquires many of its characteristics through fusion with host endosomal vesicles. Endosomal maturation is regulated by small GTPase Ras-associated binding (Rab) proteins, which localize to the vesicular membranes and recruit Rab-effector proteins involved in trafficking and fusion events [26]. Rab5 localizes to the early endosome membrane and recruits early endosome antigen-1 (EEA1), which, in co-ordination with SNARE proteins regulates early endosome homotypic fusion [27–30]. Early endosomes are pleomorphic tubulo-vesiclular structures functioning as the first recipient of internalized cargo, as well as sorting compartments for these cargos [31]. The early endosomal moderately acidic pH (6.1 – 6.8) aids in dissociating the ligands from their receptors [32], which may then be recycled back to the plasma membrane. As early endosomes mature to late endosomes, Rab5 is replaced by Rab7. Late endosomes have a luminal pH range of 6.0 – 4.9 [33] and migrate along microtubules to the perinuclear area, fusing with golgi-derived vesicles carrying newly synthesized proteases and hydrolases. Late endosomes finally fuse with lysosomes, which degrade the internalized cargo. Lysosomes maintain an acidic environment with pH~4.7, which activates degradative enzymes required for cargo degradation [32]. Thus, early endosomes, late endosomes, and lysosomes constitute a highly dynamic and adaptable continuum that trafficks and degrades cargo [34].

In order to understand the role of pH and endosomal fusion in CCV formation and maintenance, we examined endosomal maturation in *C. burnetii*-infected host cells. Our study surprisingly revealed that the CCV is significantly less acidic than the lysosomes of the mock-infected cells. Furthermore, we found that the *C. burnetii* T4BSS inhibits endosomal maturation and acidification, potentially by targeting Rab5 to Rab7 conversion on endosomes. Further, increased lysosomal biogenesis inhibits *C. burnetii* growth, suggesting host lysosomes are detrimental to *C. burnetii*.

## Results

### *C. burnetii* regulates CCV and endosomal pH in infected host cells

We previously observed the CCV pH in both HeLa cells and cholesterol-free mouse embryonic fibroblasts (MEF) to be pH~5.2, with increased CCV acidification to pH~4.8 leading to *C. burnetii* degradation [24]. Given this apparent narrow pH tolerance inside the host cell, we hypothesized that *C. burnetii* regulates the CCV pH at a less acidic pH than host lysosomes. To test this hypothesis, we compared the CCV pH from wild type (WT) *C. burnetii*-infected HeLa cells at 3 days post infection (dpi) to mature endosomal/lysosomal pH of mock-infected cells using a ratiometric microscopy-based method using pH-sensitive Oregon Green 488 dextran and pH-stable Alexa fluor 647 dextran [23]. Dextran is internalized through fluid-phase endocytosis and delivered to the CCV lumen through CCV-endosome fusion (Fig 1A). HeLa cells were pulsed with both dextrans for 4 h followed by a 1 h chase to allow for endosomal maturation, as newly formed endosomes mature and fuse with lysosomes in about 40 min [32]. The mean endosomal pH of mock-infected cells was determined from the entire cell area (Fig 1B). We found CCVs to be significantly less acidic (pH~5.2) compared to mature endosomes/lysosomes of mock-infected cells (pH~4.8) (Fig 1C). Further, the CCV pH was stable at pH~5.2 during a 6-day infection (Fig 1D), starting at 24 hour post infection (hpi). Taken together, these data suggest that *C. burnetii* regulates the CCV pH and maintains it in a less acidic range relative to host lysosomes.

**Fig 1.**
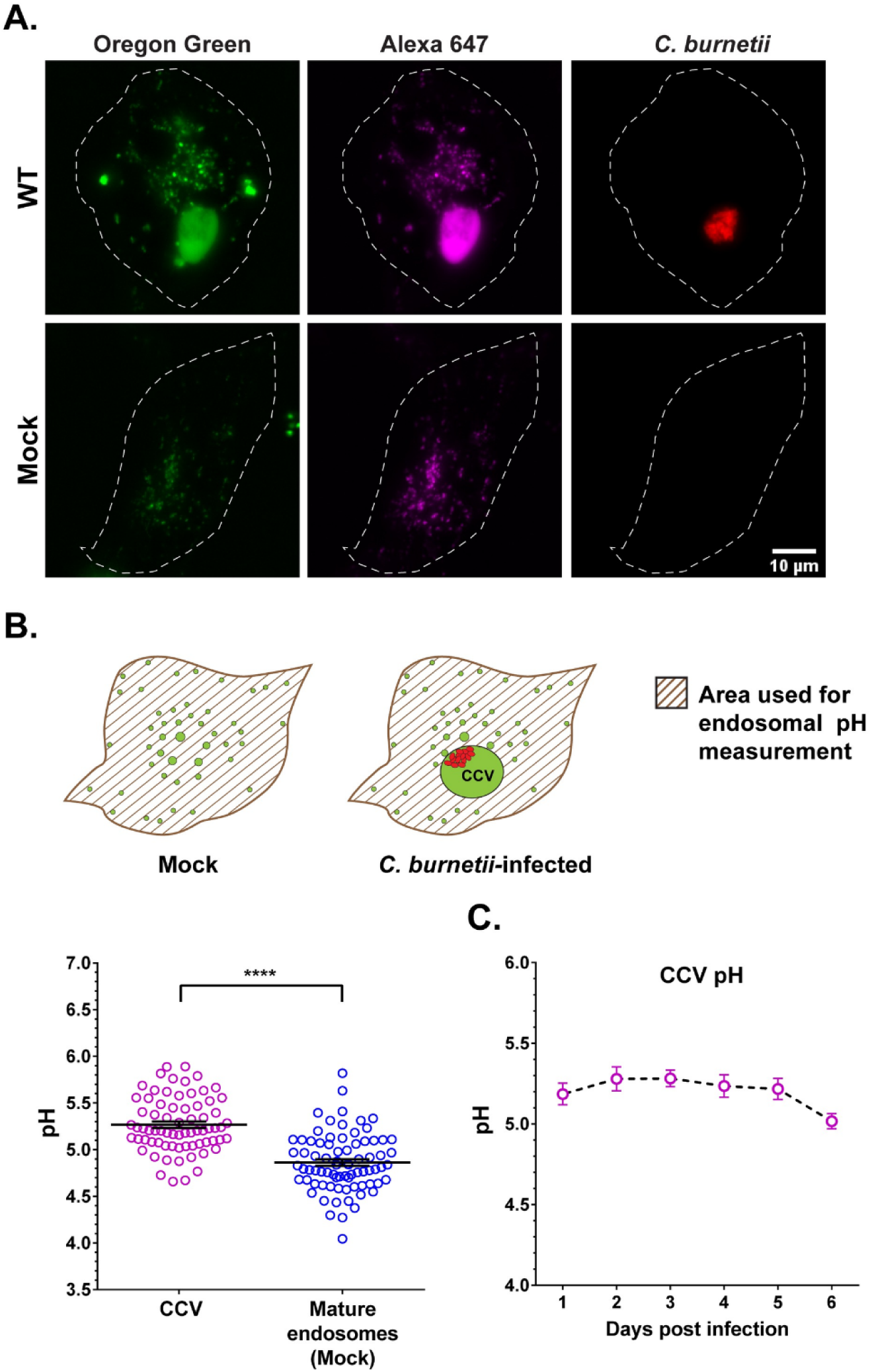
*C. burnetii* regulates CCV pH. (A) Representative images of HeLa cells infected with mCherry-*C. burnetii* and pulsed with Oregon Green 488 (pH-sensitive; green) and Alexa 647 (pH-stable; magenta) dextran for 4 h followed by a 1 h chase. Z-stacks were acquired by live cell spinning disk microscopy and Oregon Green 488 and Alexa 647 intensity quantitated. Oregon Green 488, which is quenched and less fluorescent at acidic pH, is visibly brighter in the CCV compared to the mature endosomes of mock-infected cells. (B) Diagram showing the area (patterned) used for endosomal pH measurement with the CCV excluded in infected cells. The endosomal pH is expressed as the average pH of all endosomes in the patterned area. Ratiometric pH measurement at 3 dpi revealed CCVs are significantly less acidic than the mature endosomes of mock-infected cells. Data shown as mean±standard error of mean (SEM) of at least 20 CCVs or cells in each of three independent experiments as analyzed by unpaired student t-test; ****, P<0.0001. (C) Mean CCV pH from 1 through 6 dpi revealed that the CCV pH is relatively stable at pH~5.2 without any significant difference between the days, as analyzed by one-way ANOVA with Tukey’s posthoc test. Data showed as mean±SEM of at least 15 CCVs in each of three independent experiments.

As the CCV is highly fusogenic and acquires lysosomal characteristics at least in part by fusing with host lysosomes, we examined endosomal pH in mock and WT *C. burnetii-*infected HeLa cells. By live cell microscopy, both mock and *C. burnetii-*infected cells had similar fluorescent intensity of pH-stable Alexa fluor 647, indicating equivalent internalization of fluorescent dextran. However, pH-sensitive Oregon Green 488 was visibly brighter in the “mature” endosomes in infected cells, indicating that the mature endosomes are less acidic in the infected cells (Fig 2A). Indeed, pH measurement revealed that endosomes in infected cells were significantly less acidic, with a pH~5.8 compared to those in mock-infected cells (pH~4.8) (Fig 2B). To determine if this is a bacterial-driven process, we measured mature endosomal pH in mock, WT, and Δ*dotA* (T4BSS mutant) *C. burnetii*-infected HeLa cells at 1, 2, and 3 dpi. While mature endosomes in mock and Δ*dotA*-infected cells maintained a pH ≤ 5.0, those in WT-infected cells were significantly less acidic (pH~5.8) starting 2 dpi (Fig 2C), suggesting *C. burnetii* T4BSS actively manipulates endosomal pH.

**Fig 2.**
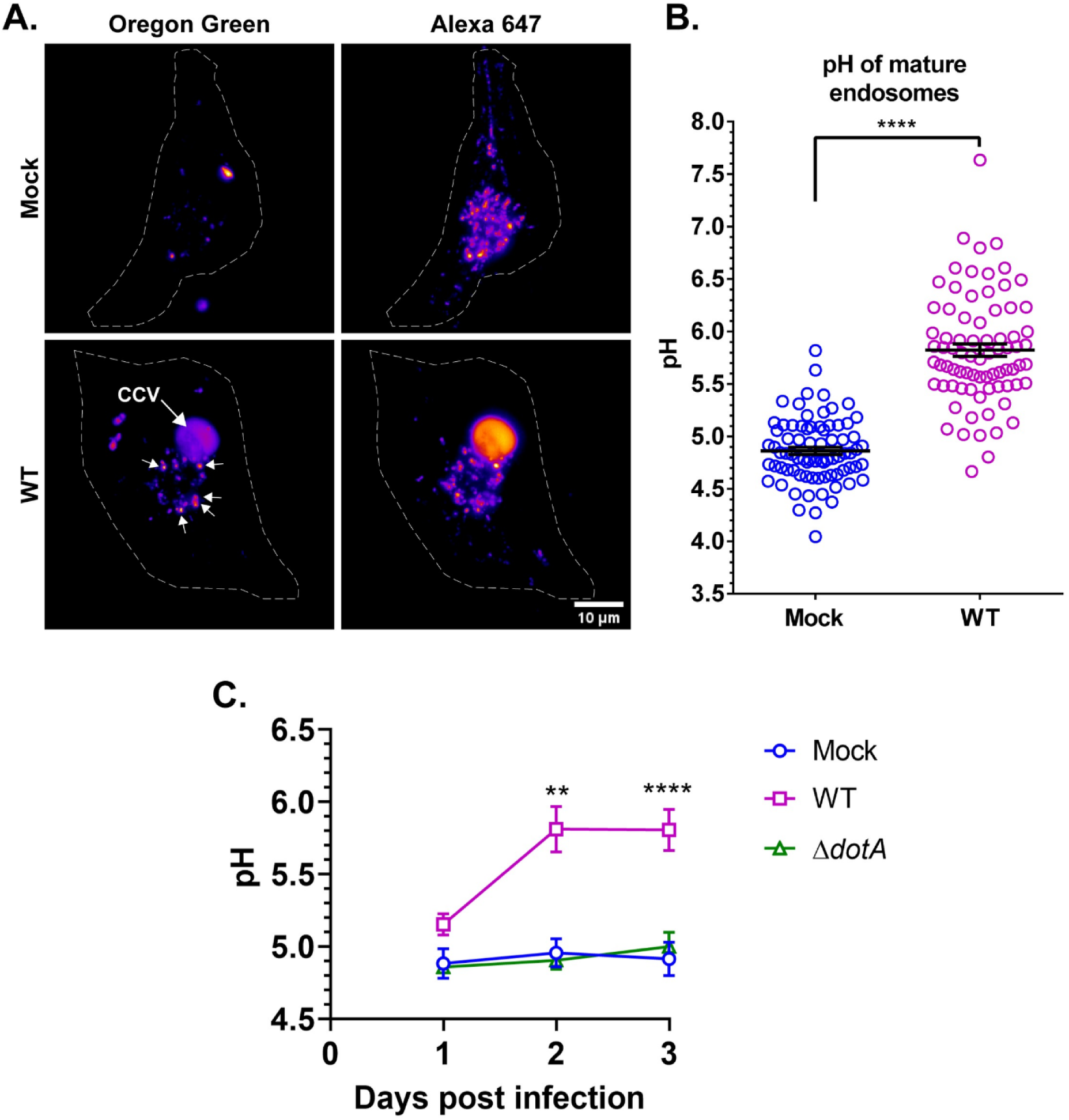
*C. burnetii* T4BSS manipulates endosomal pH in infected cells. (A) mCherry *C. burnetii-*infected HeLa cells were labeled with Oregon Green 488 and Alexa 647 dextran for 4 h followed by a 1 h chase. Z-stacked images were processed identically in ImageJ and Oregon Green 488 and Alexa 647 intensity quantitated. A heat map of mock and WT *C. burnetii*-infected cells revealed that the endosomal Alexa 647 intensity was comparable between mock and infected cells. However, Oregon Green 488 was visibly brighter in the endosomes of the infected cells (arrows), suggesting that endosomes in infected cells are less acidic. (B) Ratiometric pH measurement revealed the pH of “mature” endosomes in WT *C. burnetii*-infected cells is ~ pH 5.8, compared to ~ pH 4.9 in mock-infected cells. Data showed as mean±SEM of at least 20 cells in each of three independent experiments as determined by unpaired student t-test; ***, P<0.001. (C) Mean endosomal pH in mock, WT, and Δ*dotA*-infected cells at 1 through 3 dpi revealed that the *C. burnetii* alters endosomal maturation beginning at 2 dpi. Data shown as mean±SEM of at least 15 cells per condition in each of three independent experiments as determined by one-way ANOVA with Tukey’s posthoc test; **, P<0.01; ****, P<0.0001.

### *C. burnetii* T4BSS reduces lysosomal content in infected cells

Endosomes are progressively acidified as they mature from early endosomes (pH 6.1-6.8) to lysosomes (pH ~4.5) [33]. Therefore, the more alkaline endosomal pH we observed in WT-infected cells indicates incomplete endosomal maturation, leading us to hypothesize that the less-acidic vesicles in the WT *C. burnetii*-infected cells are not lysosomes. To test this hypothesis, we quantitated the endosomal content of mock, WT, and Δ*dotA C. burnetii-*infected HeLa cell using the early endosomal marker EEA1 (early endosome antigen 1) and the lysosomal marker LAMP1. A late endosomal marker such as Rab7 or CD63 was not included, as these proteins are also found on lysosomes, making it difficult to distinguish late endosomes from lysosomes [35, 36]. EEA1 and LAMP1 fluorescence intensities from fixed-cell microscopy images were normalized to cell area. While the entire cell was measured for mock-infected cells, the CCV was excluded for WT and Δ*dotA*-infected cells. LAMP1 intensity was reduced by 57% in WT-infected cells compared to both mock and Δ*dotA*-infected cells beginning at 3 dpi, while EEA1 intensity was unaffected at all time points (Fig 3A, B). At 6 dpi, LAMP1 was also reduced by 56% in WT-infected cells compared to mock (Fig 3A, B). However, due to the reduced viability of T4BSS mutant at late time points, the Δ*dotA* mutant was not included at 6 dpi [9]. These data suggest that the *C. burnetii* T4BSS reduces lysosomal content in infected host cells starting 3 dpi, and that the *C. burnetii* T4BSS inhibits endosomal maturation.

**Fig 3.**
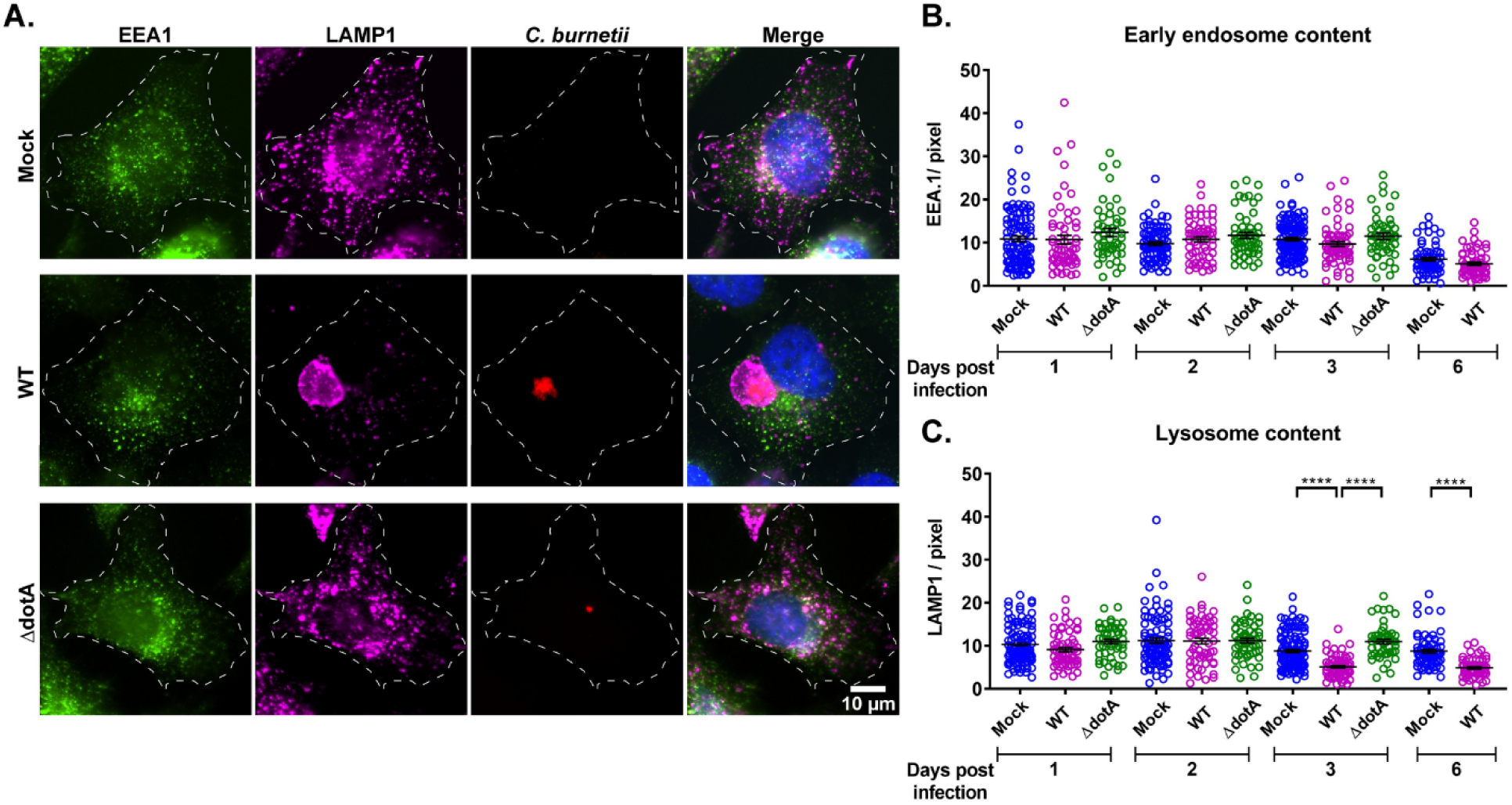
*C. burnetii* T4BSS reduces lysosomal content. (A) Representative images of EEA1 (early endosome marker) and LAMP1 (lysosome marker) immunofluorescent staining in mock and infected HeLa cells. Mock, WT, and Δ*dotA*-infected HeLa cells were immunostained with anti-EEA1, anti-LAMP1, and anti-*C. burnetii* antibodies. (B, C) Quantitation of EEA1 and LAMP1 intensity, normalized to cell area, revealed significant reduction in LAMP1 in WT *C. burnetii*-infected cells starting 3 dpi. Each circle represents an individual cell. Data shown as mean±SEM of at least 20 cells per condition in each of three independent experiments as analyzed by one-way ANOVA with Tukey’s posthoc test; ****, P<0.0001.

As discussed earlier, the CCV is highly fusogenic and undergoes homotypic and heterotypic fusion with endosomes and lysosomes during expansion between 2 and 3 dpi [37]. Thus, it is possible that the decrease in lysosomal content between 2 and 3 dpi is due to increasing CCV-lysosome fusion. To address this possibility, we measured CCV fusogenicity using a quantitative dextran trafficking assay [38]. HeLa cells infected with WT mCherry-*C. burnetii* were pulsed with fluorescent Alexa fluor 488 dextran for 10 min, followed by imaging every 4 min for 28 minutes using live-cell confocal microscopy (Fig 4A). The dextran accumulation in the CCV lumen was quantitated by measuring fluorescence intensity at every time point, and expressed as fold change over the initial time point. There was no significant difference in dextran accumulation between CCVs at 2 and 3 dpi, with an average of 2.06-fold and 1.91-fold dextran accumulation respectively (Fig 4B), suggesting that CCV fusogenicity does not change between 2 and 3 dpi. These data suggest that the reduction in lysosomes in WT *C. burnetii*-infected cells from 2 to 3 dpi is not due to increased CCV fusogenicity.

**Fig 4.**
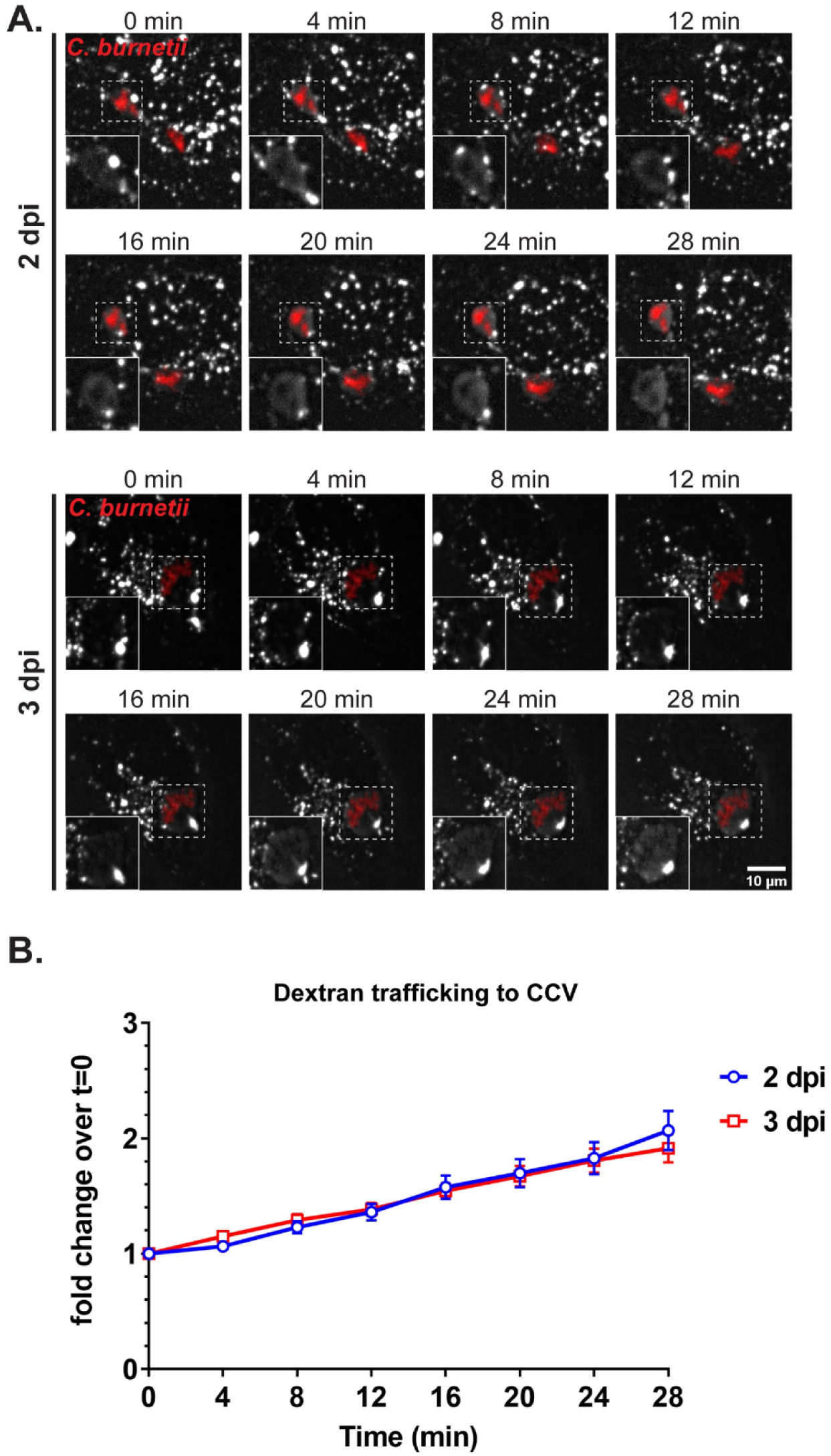
CCV fusogenicity does not increase from 2 to 3 days post infection. (A) Fluorescent dextran trafficking to CCV at 2 and 3 dpi. mCherry WT *C. burnetii*-infected HeLa cells were pulsed with Alexa 488 dextran for 10 min followed by live-cell spinning disc confocal microscopy, where the cells were imaged at 0 min post-pulse, and then every 4 min for 28 min. Images were processed identically in ImageJ; white=dextran, red=*C. burnetii*; Dextran in CCVs (boxed) shown in insets. (B) Quantification of dextran intensity in CCV revealed no significant difference in dextran trafficking to the CCV between 2 and 3 dpi. Fluorescent intensity of Alexa 488 dextran was measured from an identical region of interest (ROI) within the CCV at each time point. The mean fold change of fluorescent intensity over initial time point (t=0) was plotted against time. Data shown as the mean±SEM of fluorescent intensity fold change from at least 15 CCVs per condition in each of three independent experiments as analyzed by multiple t-test.

### *C. burnetii* T4BSS inhibits progressive endosomal acidification

Endosomes migrate along microtubules from the cell periphery towards the nucleus as they mature to lysosomes [32, 39, 40]. Indeed, peripheral endosomes are significantly less acidic than those in the perinuclear area, with “immature” endosomes mostly residing near the cell periphery whereas lysosomes aggregate in the perinuclear area [40]. To further examine *C. burnetii* regulation of endosomal maturation, we measured changes in endosomal pH from the cell periphery to the perinuclear area in mock, WT, and Δ*dotA-*infected HeLa cells. mCherry-*C. burnetii* infected HeLa cells were labelled with Oregon Green 488 and Alexa fluor 647 dextran as above, followed by a 1 h chase to allow for endosomal maturation. Images from live-cell microscopy were processed in ImageJ and cells were divided into four concentric areas (shells 1 through 3 and core as previously described in [40]; Fig 5A). For *C. burnetii*-infected cells, the CCV was excluded from each shell (Fig 5A). As expected, peripheral vesicles (shell 1) in mock-infected cells were significantly less acidic than those of the perinuclear area (core) (Table 1 & Fig 5B, C), and vesicles showed a progressive acidification from peripheral to perinuclear area. In the WT-infected cells, although the peripheral vesicles had a mean pH comparable to the mock, the perinuclear vesicles were significantly less acidic than those of mock and Δ*dotA*-infected cells at both 2 and 3 dpi (Table 1 & Fig 5B, C). These data indicate that *C. burnetii* inhibits progressive endosomal acidification in a T4BSS dependent manner.

**Fig 5.**
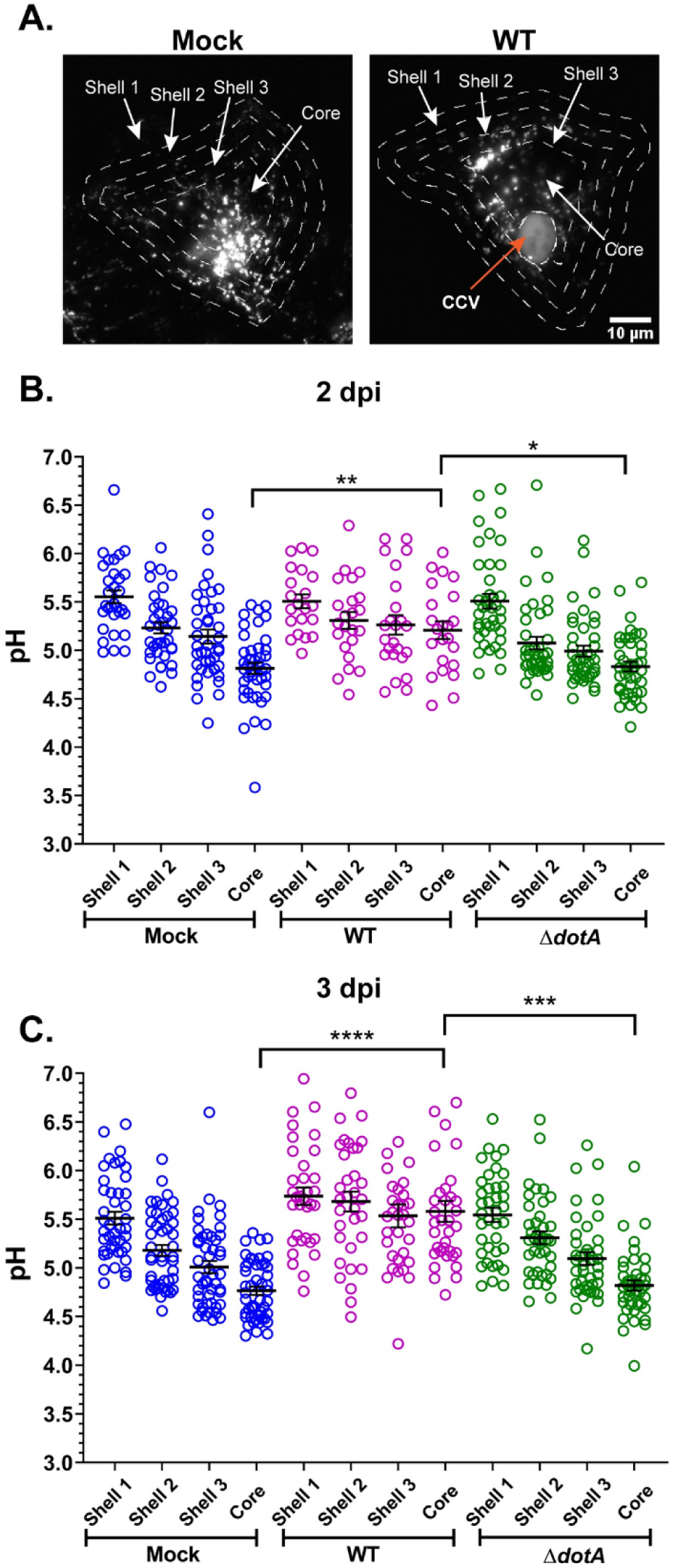
*C. burnetii* T4BSS inhibits progressive endosomal acidification. (A) Mock, mCherry WT and mCherry *ΔdotA*-infected Hela cells were pulsed with Oregon Green 488 and Alexa 647 dextran for 4 h followed by a 1 h chase for endosomal maturation. Live-cell spinning-disc microscopy images were processed identically in ImageJ. To generate concentric shells corresponding to peripheral, perinuclear, and intermediate zones, an ROI at the cell periphery was degraded 3 times, 4 um each time. In *C. burnetii*-infected cells the CCV was excluded from measurements. (B, C) Ratiometric pH analysis revealed that the mature endosomes at the perinuclear area (core) of the WT-infected cells were significantly less acidic compared to those in the mock and *ΔdotA*-infected cells at both 2 and 3 dpi. Each circle represents an individual cell. Data shown as mean±SEM of at least 15 cells per condition in each of three independent experiments as analyzed by one-way ANOVA with Tukey’s posthoc test; ****, P<0.0001; ***, P<0.001; **, P<0.01; *, P<0.05.

**Table 1.** Mean pH (± standard error of mean) of endosomes in concentric shells of mock, WT, and Δ*dotA C. burnetii*-infected HeLa cells.

| Days post infection | Area of cell | pH |  |  |
| --- | --- | --- | --- | --- |
| | | Mock | WT | $\Delta dotA$ |
| 2 | Shell 1 | 5.55±0.06 | 5.51±0.08 | 5.50±0.09 |
| | Shell 2 | 5.23±0.05 | 5.31±0.08 | 5.07±0.06 |
| | Shell 3 | 5.14±0.07 | 5.26±0.09 | 4.99±0.05 |
| | Core | 4.81±0.05 | 5.20±0.09 | 4.83±0.05 |
| 3 | Shell 1 | 5.51±0.06 | 5.73±0.08 | 5.54±0.07 |
| | Shell 2 | 5.18±0.05 | 5.68±0.10 | 5.30±0.06 |
| | Shell 3 | 5.01±0.06 | 5.53±0.11 | 5.09±0.06 |
| | Core | 4.76±0.04 | 5.58±0.10 | 4.81±0.05 |

### *C. burnetii* T4BSS dysregulates Rab5 to Rab7 conversion

The Rab GTPases Rab5 and Rab7 are key markers of early and late endosomes/lysosomes, respectively [32, 35, 36, 41]. Rab5 regulates fusion of newly formed vesicles carrying cargo with pre-existing early endosomes [42], followed by maturation to late endosomes with Rab5 being replaced by Rab7 [43]. Because Rab5 to Rab7 conversion is critical for endosomal maturation, we tested whether *C. burnetii* T4BSS affects Rab5 to Rab7 conversion using co-localization with fluorescent beads [44]. Due to the reduced phagocytic capacity of HeLa cells, we used murine alveolar macrophages (MH-S). Rab5 and Rab7 localization on phagocytosed latex beads was compared between mock, WT, and *ΔdotA-*infected cells at 15 min time intervals in a 60-min incubation. As expected, at 0 min post pulse nearly 95% bead phagosomes were positive for Rab5 in mock, WT, and *ΔdotA-*infected cells. Both mock and ΔdotA*-*infected cells progressively lost Rab5 and gained Rab7 over 60 min, with ~60% of bead phagosomes being Rab5-positive and 94% Rab7-positive (Table 2, Fig 6). In contrast, after 60 min, bead phagosomes in WT-infected cells had significantly reduced co-localization of both Rab5 and Rab7, at 41% and 68%, respectively. This data suggests the *C. burnetii* T4BSS causes rapid loss of Rab5 from phagosomes while also reducing the rate of Rab7 acquisition, giving rise to a Rab5-negative, Rab7-negative endosomal population in the host cells. While the characteristics of these Rab5-negative, Rab7-negative endosomes are unclear and warrant further investigation, these data suggest that *C. burnetii* T4BSS dysregualtes Rab conversion during endosomal maturation.

**Fig 6.**
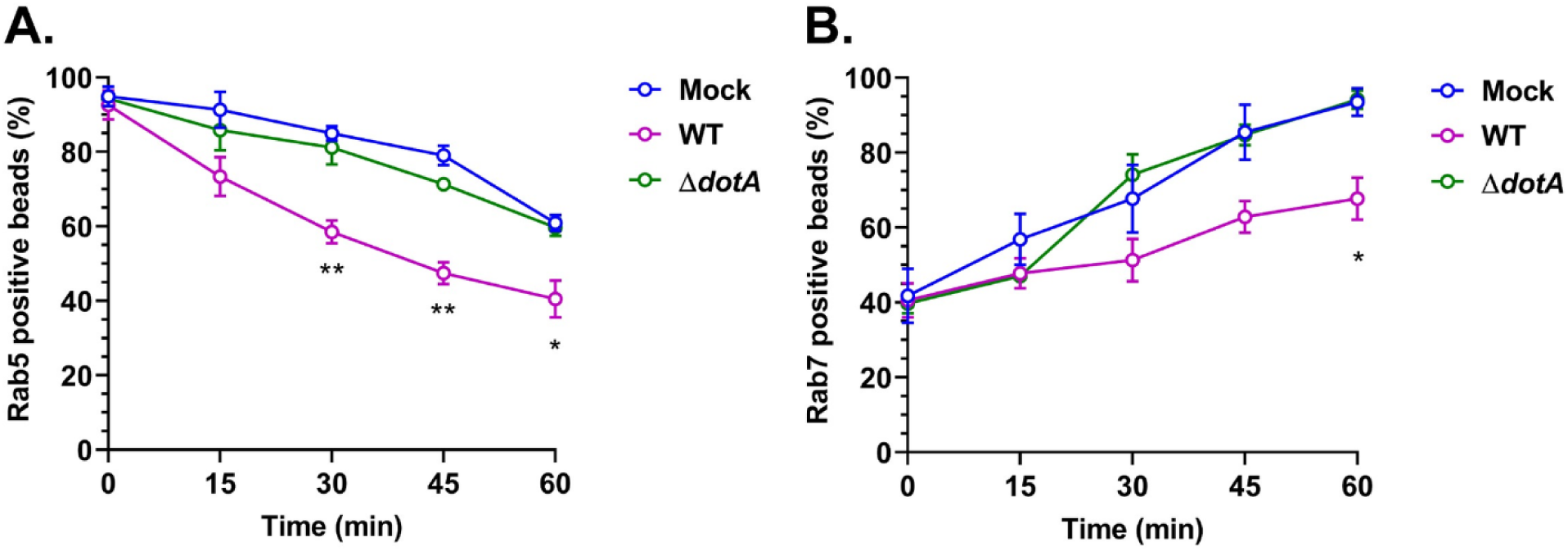
*C. burnetii* T4BSS dysregulates Rab5 to Rab7 conversion. Mock and WT *C. burnetii-*infected murine alveolar macrophages were pulsed with red fluorescent latex beads (1 um) for 15 min followed by fixation at 0 min post-pulse and every 15 min for 60 min. Cells were then immunostained for Rab5 or Rab7 and internalized beads scored for co-localization with Rab5 or Rab7. The percent of Rab5 or Rab7 positive bead phagosomes over total number of internalized beads was plotted against time. In WT-infected cells, Rab5 was lost from bead phagosome significantly faster than mock and *ΔdotA*-infected cells (A), whereas Rab7 localization to bead phagosomes was significantly slower in WT-infected cells (B). Data shown as mean±SEM of at least 30 bead phagosomes per condition in each of three independent experiments as analyzed by multiple t-test; **, P<0.01; *, P<.05.

**Table 2.**
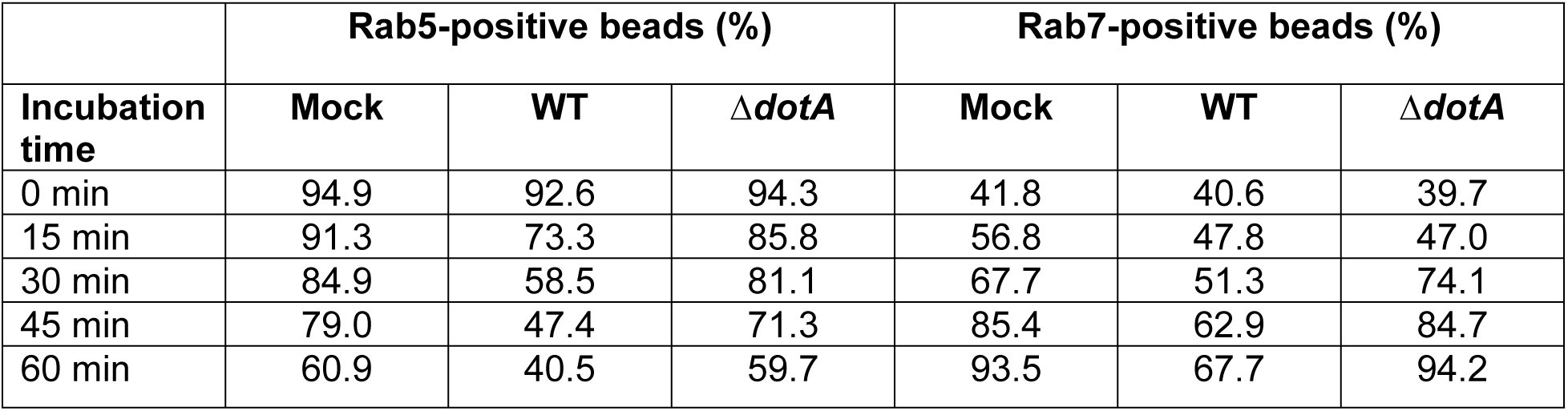
Percentage of Rab5 or Rab7 positive bead phagosomes in mock, WT, and Δ*dotA*-infected MH-S cells during a 60-min incubation.

### TFEB-induced lysosome biogenesis inhibits *C. burnetii* growth

Given that the *C. burnetii* T4BSS actively inhibits endosomal maturation and acidification to reduce lysosome content, we hypothesized that lysosomes are detrimental to *C. burnetii* intracellular growth. Lysosomal biogenesis is regulated by the transcription factor EB (TFEB), which coordinates expression of genes encoding lysosomal enzymes such as proteases and hydrolases by binding the CLEAR element in the promoter region of these genes [45]. TFEB overexpression leads to increased lysosomal biogenesis [45]. To examine whether increased lysosome biogenesis is toxic to *C. burnetii*, we measured *C. burnetii* growth in HeLa cells overexpressing TFEB-GFP [46]. First, we examined lysosomal content of the parental and TFEB-GFP HeLa cells by staining with LysoTracker an acidotropic fluorophore used for labeling acidic organelles such as lysosomes. Confirming previous observations, we found the mean LysoTracker intensity in TFEB-GFP cells to be ~2-fold higher compared to the parental cells (Fig 7A, B). Next, we analyzed *C. burnetii* growth by both immunofluorescence and quantitative colony forming unit (CFU) growth assay. Cells infected with WT *C. burnetii* were either fixed and stained with anti-*C. burnetii* antibody, or processed for CFU assay. Immunofluorescence showed fewer and smaller CCVs in TFEB-GFP cells (Fig 7C). Growth assay revealed a 50% and 70% reduction in *C. burnetii* growth at 4 and 6 dpi, respectively, in TFEB-GFP cells compared to parental cells (Fig 7D), indicating that TFEB overexpression attenuates *C. burnetii* growth. These data suggest that TFEB-induced lysosome biogenesis negatively affects *C. burnetii* growth.

**Fig 7.**
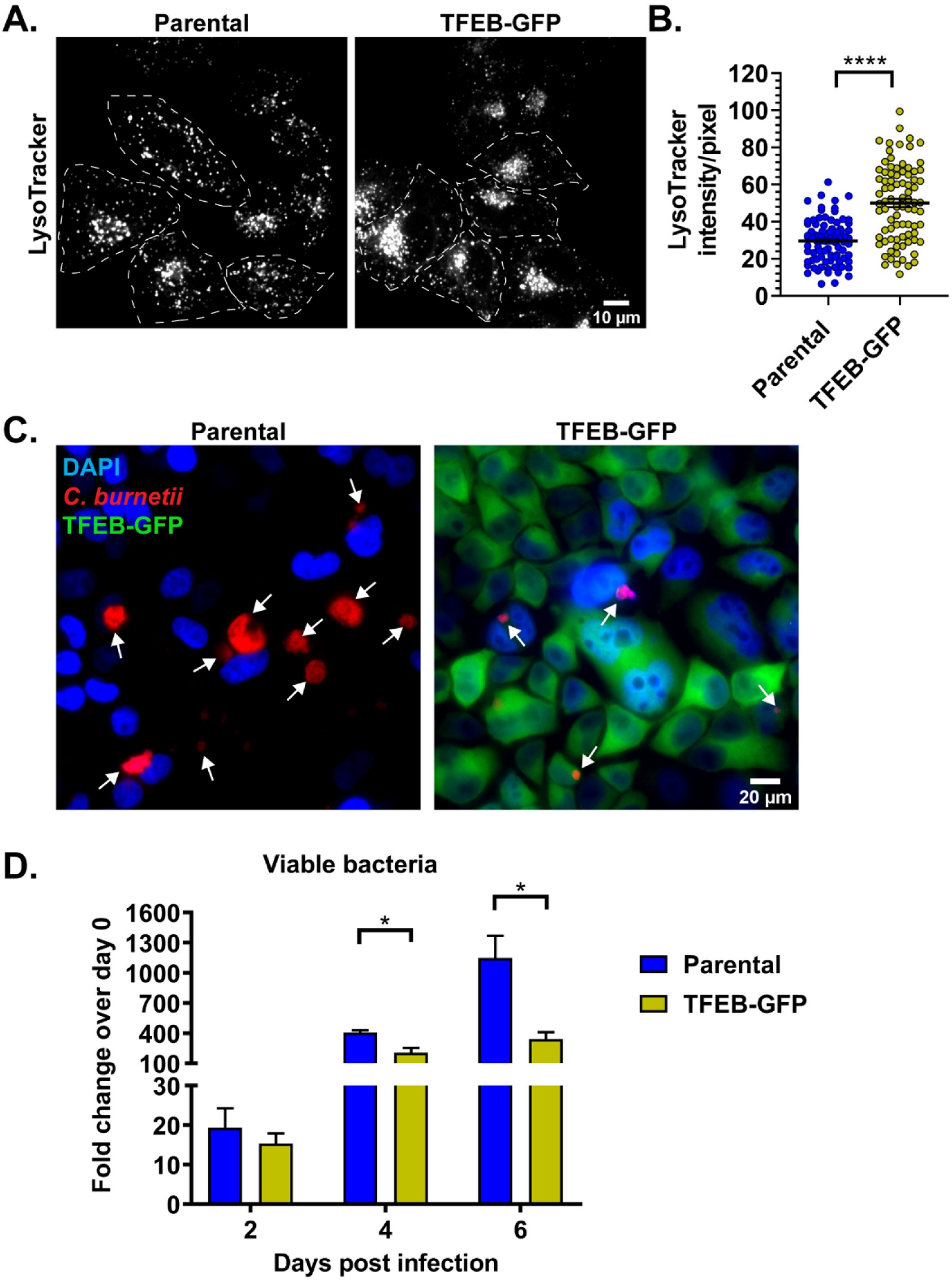
TFEB-induced lysosome biogenesis inhibits *C. burnetii* growth. (A) Representative images of parental and TFEB-GFP HeLa cells stained for 30 min with LysoTracker Red, an acidotropic fluorescent probe used to label lysosomes. Live-cell confocal images were processed identically in ImageJ. (B) Quantitation of mean LysoTracker intensity revealed a two-fold increase in LysoTracker vesicles in TFEB-GFP cells compared to parental cells. Data shown as mean±SEM of at least 25 cells per condition in each of three independent experiments as analyzed by unpaired student t-test; ****, P<0.0001. (C) Immunofluorescent staining of WT *C. burnetii*-infected parental and TFEB-GFP HeLa cells with anti–*C. burnetii* antibody. Arrows point to individual CCVs. Qualitatively, TFEB-GFP cells contained smaller and fewer CCVs compared to the parental cells. (D) Quantitative CFU assay revealed a significant reduction in bacterial growth in TFEB-GFP cells compared to parental cells. *C. burnetii* growth is plotted as a fold change of CFU/mL over 0 dpi at each time point. Data shown as mean±SEM for each condition in each of three independent experiments as analyzed by multiple t-test; *, P<.05.

## Discussion

*C. burnetii* metabolism and T4BSS secretion requires acidification of the nascent phagosome [16, 47]. However, in a previous study with cholesterol-free MEFs, we discovered the mature CCV is less acidic than previously reported (pH~5.2 versus pH~4.5) [14, 15]. Further, CCV acidification to pH<4.8 caused *C. burnetii* lysis within the CCVs, suggesting *C. burnetii* has a narrow CCV pH tolerance [24]. Here, we found that the CCV is indeed less acidic than lysosomes of mock-infected cells, further suggesting that *C. burnetii* regulates CCV pH for optimal growth. Surprisingly, *C. burnetii* inhibited progressive endosomal acidification in a T4BSS-dependent manner, leading to decreased lysosomal content and increased availability of less acidic endosomes to fuse with the CCV. Endosomes in the WT but not T4BSS mutant-infected cells underwent rapid loss of Rab5 without gaining Rab7 during maturation, suggesting the *C. burnetii* T4BSS perturbs Rab conversion to impair early to late endosome maturation. Finally, inducing lysosome biogenesis by TFEB overexpression inhibited *C. burnetii* growth, providing evidence that host lysosomes are in fact detrimental to *C. burnetii*. Together these data suggest that *C. burnetii* blocks endosomal maturation to generate a permissive replicative niche.

Many intracellular pathogens such as *Legionella pneumophila* [18], *Mycobacterium tuberculosis* [19], *Toxoplasma gondii* [48], *Chlamydia psittaci* [49], *Anaplasma sp*. [20, 21], and *Yersinia pestis* [22] block phagosome-lysosome fusion to avoid phagosomal acidification. *C. burnetii*, on the other hand, requires acidification of the nascent phagosome to activate bacterial metabolism and the T4BSS. However, we found that, beginning at 24 hours post infection, the CCV is significantly less acidic than phagolysosomes. Further, CCV pH is maintained at ~5.2 during a six day infection, indicating that *C. burnetii* regulates CCV pH. Lysosomal enzymes such as proteases and cathepsins are most active against their substrates in the pH range 3-5 [50, 51] and optimally active at pH 4.5 [50]. While the CCV is known to be proteolytically active [5, 6], the mature CCV may not be optimal for lysosomal enzyme activity, thus allowing *C. burnetii* growth. In contrast, the fact that *C. burnetii* grows well in axenic media at pH 4.75 [52] is most likely because host cell proteases and hydrolases, which are not in the media but in the CCV, are in fact detrimental to *C. burnetii*.

While the mechanism by which *C. burnetii* regulates CCV pH is not clear, *C. burnetii*-driven inhibition of endosomal maturation may indirectly affect CCV pH. Given that the CCV extensively interacts with the host endocytic pathway and acquires it’s luminal characteristics in part by fusion with endosomes and lysosomes, CCV fusion with less acidic endosomes may contribute to the decreased CCV acidity. However, additional bacterial-driven mechanisms to regulate CCV pH include inhibiting endosomal proton pumps, such as vaculolar ATPase (v-ATPse), or secreting neutralizing enzymes into the CCV lumen. The *L. pneumophila* T4SS effector protein SidK directly binds to and inhibits v-ATPase-driven proton translocation, resulting in decreased acidity of the *Legionella*-containing phagosome [53]. v-ATPase is on the CCV membrane [5], and it is possible that one or more *C. burnetii* T4BSS effectors perturb v-ATPase function as a mechanism to modulate CCV pH. *Edwardsiella ictaluri*, a channel catfish intracellular pathogen [54], replicates in a phagosome which initially acidifies to pH 4 to activate the *E. ictaluri* Type 3 Secretion System. However, *E. ictaluri* secretes an urease enzyme into the phagosome that converts urea into ammonia, reducing phagosome acidity to pH 6 [55]. *Helicobacter pylori* also survives intestinal acidity by producing urease enzyme, breaking down intestinal urea into ammonia and neutralizing intestinal acidity [56]. Whether *C. burnetii* also utilizes urease or other neutralizing factors, or modulates v-ATPase or other ion channel activity, remains to be tested.

We surprisingly found that *C. burnetii* inhibits endosomal maturation, building upon prior studies suggesting that *C. burnetii* manipulates the endocytic pathway. For example, *C. burnetii* increased transferrin accumulation in endosomes [57] and upregulated clathrin expression, a protein involved in receptor-mediated endocytosis [58, 59]. CD63, a molecular marker for late endosomes/lyososomes, was upregulated in *C. burnetii*-infected cells [57], suggesting that *C. burnetii* expands the endolysosomal compartment. While CD63 cannot differentiate between late endosomes and lysosomes, we found the average endosomal pH in *C. burnetii*-infected cells to be ~5.8, which is similar to that of late endosomes [33]. Further, *C. burnetii* significantly reduced the number of LAMP1-positive lysosomes in a T4BSS-dependent manner. Indeed, HeLa cells overexpressing TFEB, a positive regulator of lysosome biosynthesis [45] inhibited *C. burnetii* growth, suggesting that increased lysosomes are detrimental to the bacteria. Supporting this hypothesis, a recent study found that TFEB knockout macrophages promoted *C. burnetii* growth compared to wildtype macrophages [60]. Together, these studies indicate that the *C. burnetii* T4BSS expands the late endosomal compartment by blocking formation of lysosomes.

The Rab GTPases Rab5 and Rab7 are hallmarks of early and late endosomes respectively, and Rab5 to Rab7 conversion on endosomal membrane drives their maturation [43, 61, 62]. Rab5 recruitment and activation on early endosomes drives recruitment of other complexes which in turn activates and recruits Rab7, at which time early endosomes mature to late endosomes [43, 63]. Therefore, Rab5 recruitment is essential for subsequent Rab7 recruitment and endosomal maturation. We observed that the *C. burnetii* T4BSS induces Rab5 loss on bead phagosomes without subsequent Rab7 acquisition. It is possible that *C. burnetii* T4BSS induces rapid loss of Rab5, thereby preventing recruitment of other factors required for early to late endosomal maturation. Further, this may give rise to a population of Rab5-Rab7-dual negative endosomes in *C. burnetii*-infected cells. While the nature of these vesicles is not clear and requires further investigation, several intracellular bacteria are known to target Rab5-Rab7 conversion. *Listeria monocytogenes* [64] and *Tropheryma whipplei* [65] block Rab5 activation, thus arresting phagosome maturation. The *Ehrlichia chafeensis* T4SS effector Etf-2 binds active Rab5 on bead phagosomes and blocks subsequent recruitment of other proteins, effectively inhibiting endosome maturation [44]. Therefore, it is possible that the *C. burnetii* T4BSS dysregulates Rab5 to Rab7 conversion as a potential mechanism to inhibit endosome maturation.

Several *C. burnetii* T4BSS effectors are known to modulate endocytic trafficking. For example, both CvpA (CBU0665) and Cig57 (CBU1751) target the early stages of clathrin-mediated endocytosis, which is essential for *C. burnetii* growth [58, 66, 67]. CvpB (Cig2/CBU0021) localizes to early endosomes and increases CCV colocalization with the early endosome markers Rab5 and EEA1 [68]. CvpB also inhibited phosphatidylinositol-3-phosphate-5-kinase (PIKfyve), which converts early endosomal phosphatidylinositol-3-phosphate PI(3)P to phosphatidylinositol 3, 5 diphosphate (PI(3,5)P2) during endosomal maturation. PI(3)P is targeted by several other intracellular bacteria as a mechanism to block maturation of the bacteria-containing phagosome. For example, *M. tuberculosis* secretes the PI(3)P phosphate SapM, reducing PI(3)P levels and delaying phagosome maturation [69, 70]. *L. pneumophila* phospholipase VipD removes PI(3)P from phagosomal membrane, effectively arresting phagosome maturation [71]. It is possible that *C. burnetii* CvpB plays a role in blocking endosomal maturation during *C. burnetii* infection; however, the lack of a significant growth phenotype in a cvpB mutant suggests that multiple, unidentified T4BSS effector proteins are involved [68, 72].

Our data indicates that while the CCV initially matures into an acidic phagolysosome, *C. burnetii* blocks further CCV acidification and maintains the CCV pH>5 during CCV expansion and bacterial replication. *C. burnetii* T4BSS effector protein(s) induce rapid Rab5 loss on endosomes and expand the late endosomal compartment, potentially as a mechanism to indirectly modulate CCV pH and proteolytic activity. These findings suggest that *C. burnetii* is in fact sensitive to the lysosomal environment and not passively acidified, and *C. burnetii* regulation of CCV pH is a critical step in *C. burnetii* pathogenesis.

## Materials and methods

### Bacteria and mammalian cells

*Coxiella burnetii* Nine Mile Phase II (NMII) (clone 4, RSA 439), Δ*dotA* (T4BSS mutant) *C. burnetii,* and mCherry-expressing wild-type (WT) *C. burnetii* [24] were grown for 4 days in ACCM-2, washed twice with phosphate buffered saline (PBS) and stored as previously described [52]. A mCherry-expressing Δ*dotA C. burnetii* was generated by electroporating pJB-CAT-1169-mCherry into Δ*dotA C. burnetii* as described previously [73]. The multiplicity of infection of each bacteria stock was optimized for each cell type to ~1 bacterium per cell. Human cervical epithelial cells (HeLa, ATCC CCL-2) and mouse alveolar macrophages (MH-S; ATCC CRL-2019) were maintained in RPMI (Roswell Park Memorial Institute) 1640 medium (Corning, New York, NY, USA) containing 10% fetal bovine serum (FBS; Atlanta Biologicals, Norcross, GA, USA) and 2 mM L-alanyl-L-glutamine (glutagro; Cat. 25-015-CI, Corning, New York, NY) at 37°C and 5% CO_2_. The wild type (parental) and TFEB-GFP expressing HeLa cells (generously provided by Richard J. Youle) [46] were maintained in DMEM (Dulbecco’s Modified Eagle Medium; Corning) containing 10% FBS at 37°C and 5% CO_2_.

### CCV and vesicular pH measurements

The CCV pH was measured as described previously [23] with modifications. Briefly, HeLa cells were infected with mCherry-WT or mCherry-Δ*dotA C. burnetii* in six-well plates for 2 h, washed extensively with phosphate buffered saline (PBS), and incubated in 10% RPMI. On the day before the indicated time points, cells were trypsinized, resuspended to 1X10^5^ cells/mL, and plated onto ibidi-treated channel *µ*-slide VI^0.4^ (3X10^3^ cells per channel; ibidi USA Inc., Verona, WI). The next day cells were labeled with pH-sensitive Oregon Green 488 dextran (MW, 10,000; Invitrogen) and pH-stable Alexa fluor 647 dextran (MW, 10,000; Invitrogen) at a final concentration of 0.5 mg/mL in 10% RPMI for 4 h followed by a 1 h chase to allow for endosomal maturation. After washing with PBS, cells were incubated in 10% RPMI and individual CCVs imaged live using z-stacks of 0.2 µm steps with a Nikon spinning disk confocal microscope (60X oil immersion objective) and Okolab Bold Line stage-top incubator for environmental control (Okolab USA Inc., San Bruno, CA). Images were captured and processed identically, fluorescence intensity from maximum intensity projections was measured for Oregon Green 488 and Alexa fluor 647 using ImageJ (Fiji; [74]) and 488/647 ratio was calculated. For measuring lysosomal pH from mock-infected cells, 488/647 ratio of the entire cell was calculated, whereas for the same analysis from *C. burnetii-*infected cell, the CCV was excluded from the cell area. For measuring endosomal pH in concentric shells [40], first a region of interest (ROI) was drawn around the cell periphery, which was then scaled down three times, 4 µm each time to generate three concentric shells (S1 through 3) and a core (C). In *C. burnetii-*infected cells, the CCV was subtracted from each of these areas. 488/647 ratio of the areas were used to determine the mean endosomal pH. To generate a pH standard curve, wild type *C. burnetii*-infected HeLa cells (3 days post infection; dpi) were incubated in equilibration buffer (143 mM KCl, 5 mM glucose, 1 mM MgCl_2_, 1 mM CaCl_2_, and 20 mM HEPES) containing ionophores nigericin (10 uM) and monensin (10 uM) for 5 min followed by incubation in standard buffers of pH ranging from 4.0 to 7.0 containing ionophores for 5 min before imaging. At least 20 CCVs were measured at each pH and the 488/647 ratio was plotted against the pH of the respective buffer to obtain a sigmoidal standard curve. The experimental samples were then interpolated to the standard curve to determine the pH; a standard curve was generated for each individual experiment. At least 20 CCVs/cells were measured for each experimental time point per condition for three independent experiments.

### Quantitation of early endosomes and lysosomes

HeLa cells were infected with WT or Δ*dotA C. burnetii* in six-well plates (5X10^4^ cells per well; two wells for each time point) for 2 h, washed extensively with PBS, and incubated in 10% RPMI. On the day before the indicated time points, cells were trypsinized and resuspended to 1X10^5^ cells/mL, replated onto coverslips placed in 24-well plate (5X10^4^ cells per coverslip), and allowed to adhere overnight. Cells were fixed in 2.5% paraformaldehyde (Cat. 15710, Electron Microscopy Sciences; Hatfield, PA, USA) for 15 min and blocked/permeabilized for 20 min in 1% bovine serum albumin (BSA) and 0.1% saponin in PBS. Cells were then incubated in mouse anti-EEA1 (1:500; Cat. 610546; BD Biosciences, San Jose, CA), rabbit anti-LAMP1 (1:1000; Cat. ab24170 Abcam, Cambridge, MA), and guinea pig anti-*C. burnetii* (1:2500; Robert Heinzen, NIH, Hamilton, MT) for 1h followed by Alexa fluor secondary antibodies (1:1000; Life Technologies) for 1 h. Following washing with PBS, coverslips were mounted using ProLong Gold with 4’, 6’-diamidino-2 phenylindole (DAPI) (Life Technologies), and visualized on a Nikon TiE fluorescent microscope using 60X oil immersion objective. Images were captured and processed identically and the fluorescent intensity of EEA1 and LAMP1 were measured (ImageJ) and normalized to cell area. The CCV was excluded when measuring the fluorescent intensities in *C. burnetii*-infected cells. At least 20 cells were measured per condition for each of three independent experiments.

### Dextran trafficking

Dextran trafficking and fusion with CCVs was measured as described previously [38]. Briefly, HeLa cells were infected with mCherry-WT *C. burnetii* in six-well plates (5X10^4^ cells/well; two wells per condition). On the day before the indicated time points, cells were trypsinized, resuspended to 3X10^5^ cells/ml, and replated onto ibidi slides (9X10^3^ cells per channel). On a Nikon spinning disk confocal microscope (60X oil immersion objective) with Okolab Bold Line stage top incubator, CCVs were identified and marked using NIS elements (Nikon) prior to labeling with Alexa fluor 488 dextran (MW 10,000, Invitrogen) for 10 min in 10% RPMI. The cells were washed with PBS 5-6 times, and replaced with 10% RPMI. Z-stacked confocal images were obtained for each CCV every 4 min for 28 min (t=0 through 28; 8 time points). The mean dextran fluorescent intensity of an identical region of interest (ROI) within each CCV was quantified for each time point (ImageJ). The fold change of dextran fluorescent intensity over initial time point (t=0) was plotted against time. At least 15 CCVs were imaged per condition for each of three independent experiments.

### Quantification of Rab5/Rab7 positive phagosomes

MH-S cells were infected with WT or Δ*dotA C. burnetii* in six-well plates (1X10^5^ cells/well; three wells per condition). At 1 dpi, cells were trypsinized and resuspended to 1X10^5^ cells/ml, and then replated onto coverslips placed in 24-well plate (5X10^4^ cells per coverslip). Red fluorescent beads (180 µL; 1 µm FluoSpheres; Cat. F13083, Life Technologies, Eugene, OR) were centrifuged at 10,000 xg for 1 min, washed once with PBS, centrifuged, and resuspended in 9 mL 10% RPMI to 2X10^8^ beads/mL. MH-S cells on coverslips were pulsed with 250 µL (1000 beads per cell) of bead suspension for 15 min, washed with PBS, and incubated in 10% RPMI. A set of coverslips containing mock, wild type, and Δ*dotA*–infected cells were transferred to a separate 24-well plate with 2.5% PFA at 0 min post-pulse and every 15 min thereafter for 60 min. Cells were fixed for 15 min, washed in PBS, and blocked/permeabilized in 1% BSA and 0.1% saponin in PBS for 20 min. Cells were then separately stained for Rab5 or Rab7 by incubating in either rabbit anti-Rab5 (1:100; Cat. 2143S Cell Signaling Technology, Danvers, MA) or rabbit anti-Rab7 (1:100; Cat. ab137029, Abcam) along with guinea pig anti-*C. burnetii* (1:2500) for 1 h followed by Alexa fluor secondary antibodies (1:1000) for 1 h. Following washing with PBS, cells were incubated with Alexa fluor 647 conjugated wheat germ agglutinin (WGA; Cat. W32466, Life Technologies) in PBS for 30 min to stain the plasma membrane. Following washing with PBS, the coverslips were mounted using ProLong and visualized (Nikon TiE fluorescent microscope, 60X oil immersion objective). Cells were scored for internalized beads (as determined by WGA staining) and for Rab5 or Rab7 positive bead phagosomes, and colocalization with beads expressed as the percent of Rab5 or Rab7 positive beads over total number of internalized beads. At least 30 beads were counted (approximately 20 cells) per time point per condition for each of three independent experiments.

### LysoTracker Red staining

LysoTracker Red staining of parental and TFEB-GFP HeLa cells was performed according to the manufacturer’s protocol (LysoTracker Red DND-99; Cat. L7528, Life Technologies). Briefly, parental and TFEB-GFP HeLa cells were seeded in ibidi µ-slides (9X10^3^ cells per channel) and allowed to adhere overnight. Cells were labeled with 100 nM LysoTracker Red in 10% DMEM for 30 min, washed with PBS to remove uninternalized LysoTracker, and replaced with 10% DMEM. Confocal images were obtained from live cells using Nikon spinning disk confocal microscope (60X oil immersion objective) with Okolab Bold Line stage top incubator for environmental control. Images were captured and processed identically in ImageJ software. The mean LysoTracker intensity, normalized to cell area, was measured for at least 25 cells for both parental and TFEB-GFP HeLa cells in each of three independent experiments.

### *C. burnetii* viability by colony forming unit (CFU) assay

Parental and TFEB-GFP HeLa cells were plated in six-well plate (2X10^5^ cells per well) and allowed to adhere overnight. The cells were infected with wild type *C. burnetii* in 0.5 mL DMEM for 2 h, washed extensively with PBS, and scraped into 2 mL of fresh 10% DMEM. Infected cells were replated into a 24-well plate (2.5X10^4^ cells/well for day 2, 10^4^ cells/well for day 4, and 5X10^3^ cells/well for day 6). To determine day 0, 500 µL (5X10^4^) of infected cells were lysed in sterile water for 5 min. The released bacteria were diluted 1:5 in ACCM-2 and plated in 5-fold serial dilutions onto 0.25% ACCM-2 agarose plates [75]. For the subsequent time points, the cells were lysed in sterile water for 5 min and the released bacteria were diluted 1:5 in ACCM-2 and spotted in 10-fold serial dilutions onto 0.25% ACCM-2 agarose plates. The plates were incubated for 7 to 9 days at 37°C in 2.5% O_2_ and 5% CO_2_, and the number of colonies counted to measure bacterial viability. Each of the three experiments was performed in biological duplicate, and the bacteria were spotted in triplicate.

### Data analyses

Image processing and analyses were done in ImageJ (Fiji) software [74]. Statistical analyses were performed using unpaired student t-test, ordinary one-way ANOVA (with Tukey’s correction), or multiple t-test as appropriate in Prism (GraphPad, La Jolla, CA).

## Acknowledgments

We thank Richard J. Youle (National Institute of Neurological Disorders and Stroke) for providing the parental and TFEB-GFP HeLa cells. We thank Seth Winfree for assistance with quantitative microscopy and insightful discussions. We also thank Rajendra Angara, Baleigh Schuler, Rochelle Ratnayake, and Piya D. Ghatak for critical reading of the manuscript and the IUSM Biology of Intracellular Pathogens Group for helpful suggestions.

